# Population ecology of *Trigoniulus corallinus* (Gervais) (Diplopoda: Spirobolida)

**DOI:** 10.1101/2020.08.23.263855

**Authors:** S. Bhakat

## Abstract

A population of *Trigoniulus corallinus* (Gervis) in an open land rich in organic matter is studied for a year. Population density and biomass ranged from 2.13 to 56.31per m^2^ and 1026.38 to 8494.38 per m^2^ respectively. Various indices showed that population of *T. corallinus* is aggregated in distribution in the peak period of their abundance and this is due to patchy distribution of food, soil moisture and sexual attraction. Monthly age structure showed adult and late stadia are abundant in June, October and November while August population covered all the stadia.

In the developmental stages, length and width progression factor of *T. corallinus* ranged from 1.11 to 1.98 (mean 1.43) and 1.10 to 1.56 (mean 1.31) respectively. Weight progression factor in female is higher compared to that of male and this may due to more accumulation of egg forming tissue in female. Population density and biomass is significantly correlated with minimum temperature and rainfall.

## Introduction

*Trigoniulus corallinus* (Gervais), a senior synonym of *Trigoniulus lumbricinus* (Gerstaecker) (Shelley and Lehtinen, 1999) is a very common millipede of West Bengal (Mukherjee, 1962; Bhakat, 2014). It is mainly found in garden soil which is rich in plant debris and other organic matters. So far no work on its population ecology have been made on this millipede except the study of Banerjee (1974) who reported annual fluctuation in population and correlated population density with rainfall in north-east India. Mukherjee (1962) recorded its variation in segment number from West Bengal. As the millipede is a tropical species, there are several reports on its occurrence in other states of India (Chezhian and Prabakaran, 2016; Choudhari et al. 2014; Shukla and Tripathi, 1974). Most of the works are restricted to physiology of *T. lumbricinus* (Tripathi and Shukla, 1980; Shukla, 1982, 1984a, 1984b; Shukla and Misra, 1984; Shukla and Shukla 1981, 1986; Bhakat, 2014). So it became necessary to study the ecology of *T. corallinus* in detail. In this context, the present paper describes the population density, biomass, dispersion pattern and seasonal age structure of *T. corallinus* as observed in an open ecosystem.

## Materials and methods

Samples were collected regularly from Nalhati (87°47’E, 24°19’N) (District Birbhum, West Bengal). The sampling site at Nalhati was an open land (3600 sq.m.) rich in organic matter (as it is used as dumping ground of house hold wastes like skin of vegetables, rotten vegetables, kitchen waste etc. and faecal pellets of domestic animals like sheep, goat). During the study period (June 2008 to May 2009) seasonal maximum, minimum temperature and rainfall data were recorded and presented in Table1.

The millipede population was sampled using a folding quadrat of one square meter (Bhakat, 1987). The animals were first picked up of the surface, the leaf litters and other contents of the sample then placed in a plastic jar for searching in the laboratory. The underground members (upto a depth of 15 cm) were collected separately. Sampling sites were randomly selected. Samples were made at weekly intervals and in each week four quadrats were taken.

Dispersion pattern of *T. corallinus* was measured by using different indices followed by Bhakat (1989). Aggregation has been measured by using k in the expression S^2^ = m + km^2^, derived from the negative binomial distribution (Bliss and Owen, 1958), where, S^2^ is the variance and m is the mean. In this method the degree of aggregation is determined from the value of k, which in case of an extreme aggregation approaches zero. The mean crowding (m*) is defined by Lloyd (1967) as the mean number of other individuals per individual per quadrat. It relates to the mean population density (m) and the variance (S^2^) of the population in the following way: m* = m + (S^2^/m – 1). The ratio of mean crowding to mean density m*/m, is called patchiness by Lolyd, which is identical with Morisita’s (1971) dispersion index I*_A_ and Kuno’s (1968) index C_A_. m*/m = (m^2^ + S^2^ – m)/m = 1 + C_A_ = I*_A_. The m*/m provides a relative measure of aggregation. It equals unity in random and greater and smaller than unity in aggregated and regular distributions respectively. A X^2^ test for small samples (Elliott, 1973) was used to assess distribution with the Poisson series, where X^2^ = S^2^(n – 1)/mean. Southwood (1966) further showed that index of dispersion can be measured in the following way: Index of dispersion = X^2^/(n – 1). The value approximate to unity in random distribution, a value of zero for regularly distributed and a value of greater than one indicates aggregation. Measurement for aggregation was not calculated for the months of January to April as the population sizes for these months were smaller and samples were mainly consisted of 0’s and 1’s.

Age was determined by the number of rows of ocelli (Vachon, 1947). Each moult was followed by regular addition of a row of ocelli like 2, 3, 4etc. As for example in the stadium IV there were three rows of ocelli and in first, second and third rows number of ocelli was one, two and three respectively. The rows of ocelli were parallel to each other and form a triangular shape.

Lengths and widths of different stadia were determined as described by Blower and Gabbut (1964), Blower and Miller (1977) and Bhakat (1987). For the estimation of biomass each was weighed separately with an accuracy upto 0.1 mg. Growth or weight progression factor was calculated by the following formula:

Growth or weight progression factor (P_f_) = Growth or weight of (X+1) stadium/Growth or weight of X stadium. Where, X = I, II, …..IX stadia.

## Results

### Population density and biomass

Population density and biomass of *T. corallinus* during different months of the study period is presented in Table 2. The population density varied from 2.13 (February) to 56.31 (August) per m^2^. The millipedes were found in high densities during June to October and minimum in numbers during other months of the year. Biomass values ranged from 1026.38mg (February) to 8494.38mg (June) per m^2^. Though population density was maximum during August but the biomass was maximum in the month of June as the population consisted of most adults and gravid females which were heavier.

**Table 1.** Monthly weather data of Nalhati during the study period (June, 2008 to May, 2009).

| Parameter/Month | June | July | Aug. | Sept. | Oct. | Nov. |
| --- | --- | --- | --- | --- | --- | --- |
| Max. temperature (°C) | 34.1 | 33.2 | 33.0 | 34.0 | 32.9 | 30.5 |
| Min. temperature (°C) | 23.7 | 24.1 | 23.5 | 23.1 | 19.7 | 15.5 |
| Rainfall (mm) | 320.5 | 243.4 | 307.8 | 130.5 | 177.6 | 21.5 |
| Parameter/Month | Dec. | Jan. | Feb. | Mar. | Apr. | May |
| Max. temperature (°C) | 27.5 | 24.7 | 30.0 | 33.2 | 40.6 | 38.3 |
| Min. temperature (°C) | 12.5 | 10.0 | 11.2 | 16.1 | 23.0 | 23.4 |
| Rainfall (mm) | 4.0 | - | 3.1 | 11.3 | 6.2 | 250.8 |

**Table 2.** Monthly population density and biomass of *T. corallinus*.

| Parameter | June'08 | July | Aug. | Sept. | Oct. | Nov. |
| --- | --- | --- | --- | --- | --- | --- |
| Population density m<sup>-2</sup> | 20.79 | 26.28 | 56.31 | 52.08 | 33.87 | 19.07 |
| Biomass mg.m<sup>-2</sup> | 8494.38 | 5212.20 | 6549.21 | 5390.37 | 5605.84 | 5605.94 |
| Parameter | Dec. | Jan.'09 | Feb. | March | Apr. | May |
| Population density m<sup>-2</sup> | 10.85 | 3.09 | 2.13 | 3.97 | 8.85 | 10.58 |
| Biomass mg.m<sup>-2</sup> | 4208.14 | 1285.93 | 1026.38 | 1342.65 | 2722.56 | 2835.98 |

### Dispersion pattern

The values of different indices for different months are shown in Table 3. All the values show that *T. corallinus* are aggregated in distribution in the peak period of their abundance.

**Table 3.**
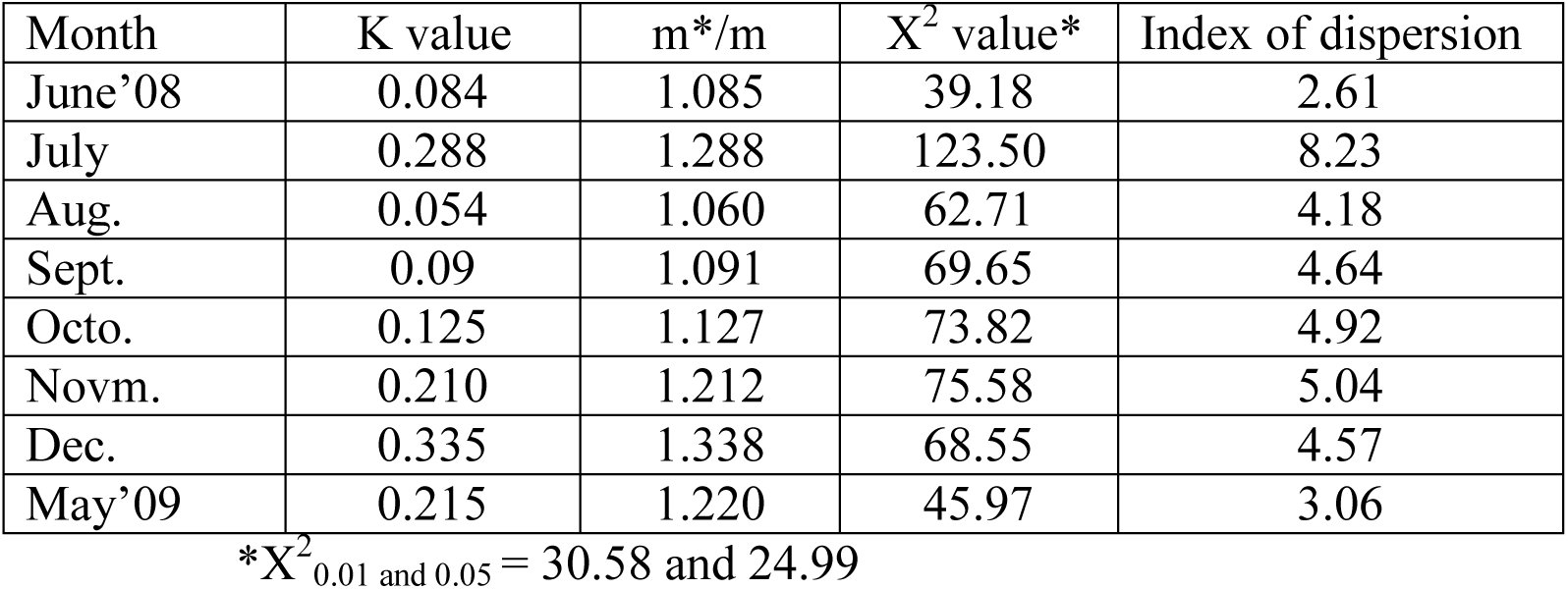
Different indices used to show dispersion pattern of *T. corallinus*.

| Month | K value | m*/m | X<sup>2</sup> value* | Index of dispersion |
| --- | --- | --- | --- | --- |
| June'08 | 0.084 | 1.085 | 39.18 | 2.61 |
| July | 0.288 | 1.288 | 123.50 | 8.23 |
| Aug. | 0.054 | 1.060 | 62.71 | 4.18 |
| Sept. | 0.09 | 1.091 | 69.65 | 4.64 |
| Octo. | 0.125 | 1.127 | 73.82 | 4.92 |
| Novm. | 0.210 | 1.212 | 75.58 | 5.04 |
| Dec. | 0.335 | 1.338 | 68.55 | 4.57 |
| May'09 | 0.215 | 1.220 | 45.97 | 3.06 |

\*X<sup>2</sup><sub>0.01 and 0.05</sub> = 30.58 and 24.99

### Age structure

There are nine stadia in the life cycle of *T. corallinus*. Data on segment numbers and number of podous and apodous segments in each stadium are presented in Table 4. The distribution of stadia in different months of the year is shown in Table 5. In June 50% of the population was adult and among immature, majority was stadium VIII. July population consisted of various stadia except stadium V while August population covered all the stadia. In September, middle stadia i. e. stadia IV, V and VI were maximum in number. In October and November late stadia were high in density. In December the stadial structure was very similar to that of June but the population size was smaller. From January’09 to March’09 the sample sizes were small and in these months only adults (stadium IX) and stadium VIII were found in maximum number. Then population sizes steadily increased in April and May and mostly consisted of stadia VII, VIII and adults.

**Table 4.** Number of podous and apodous segments in different stadium of *T. corallinus*.

| Stadium | No. of podous segment (mean) | No. of apodous segment (mean) | N |
| --- | --- | --- | --- |
| II | 8 | 4 | 19 |
| III | 12 | 4 | 21 |
| IV | 17 (15) | 5 | 28 |
| V, Male | 22 – 34 (23) | 5 | 32 |
| Female | 22 – 34 (23) | 5 | 19 |
| VI, Male | 29 – 37 (34) | 3 – 7 (5) | 31 |
| Female | 34 – 41 (38) | 2 – 6 (5) | 53 |
| VII, Male | 42 – 45 | 3 | 18 |
| Female | 40 – 47 (45) | 2 – 4 | 51 |
| VIII, Male | 44 – 47 | 1 – 4 | 57 |
| Female | 48 – 50 | 0 – 2 | 34 |
| IX, Male | 47 – 48 | 1 – 2 | 52 |
| Female | 50 – 52 | 0 – 1 | 49 |

**Table 5.** Stadial composition of *T. corallinus* during different months of the year (in percentage).

| Month/Stadium | II | III | IV | V | VI | VII | VIII | IX |
| --- | --- | --- | --- | --- | --- | --- | --- | --- |
| June'08 | - | - | - | - | 3.0 | 11.5 | 34.0 | 51.5 |
| July | 20.5 | 18.5 | 4.5 | - | - | 9.5 | 22.0 | 25.0 |
| August | 18.4 | 16.0 | 12.6 | 13.0 | 8.0 | 10.0 | 13.0 | 9.0 |
| September | - | 2.5 | 20.2 | 23.5 | 23.6 | 14.0 | 9.2 | 7.0 |
| October | - | - | - | 8.2 | 26.0 | 32.1 | 22.2 | 11.5 |
| November | - | - | - | 1.9 | 7.6 | 27.8 | 33.0 | 29.8 |
| December | - | - | - | - | 4.0 | 19.0 | 29.0 | 48.1 |
| January'09 | - | - | - | 1.8 | 3.8 | 3.8 | 33.2 | 57.4 |
| February | - | - | - | - | - | 5.0 | 41.4 | 53.6 |
| March | - | - | - | - | - | 6.8 | 34.6 | 58.6 |
| April | - | - | - | - | 1.9 | 13.0 | 39.6 | 45.5 |
| May | - | - | - | - | 9.2 | 20.8 | 47.0 | 23.0 |

### Development

The data on length, width, fresh weight and P_f_ of different stadia are presented in Table 6. Growth progression factor of *T. corallinus* on the basis of length and width ranged from 1.11 to 1.98 and 1.10 to 1.56 respectively. Mean value of P_f_ of male and female is not significantly different while weight progression factor of two sexes are significantly different (P< 0.01).

**Table 6.** Characteristics of different stadia and progression factor of *T. corallinus*.

| Stadium | Length, mm<br>(mean) | P<sub>f</sub> | Width, mm<br>(mean) | P<sub>f</sub> | Weight, mg<br>(mean) | P<sub>f</sub> |
| --- | --- | --- | --- | --- | --- | --- |
| II | 3.2-5.9<br>(4.21) |  |  |  |  |  |
| III | 5.7-6.8<br>(6.14) | 1.46 |  |  | 1.19-2.15 (1.82) |  |
| IV | 6.3-7.5<br>(6.85) | 1.12 | 0.90-1.10<br>(1.01) |  | 6.45-10.25 (9.62) | 5.28 |
| V, Male | 8.6-9.2<br>(8.93) | 1.30 | 1.12-1.14<br>(1.13) | 1.12 | 14.05-20.85 (18.29) | 1.90 |
| Female | 9.0-9.5<br>(9.17) | 1.34 | 1.15-1.50<br>(1.35) | 1.34 | 14.50-21.50 (18.94) | 2.07 |
| VI, Male | 14.5-22.0<br>(16.86) | 1.89 | 1.51-2.15<br>(1.71) | 1.51 | 38.74-91.75 (50.14) | 2.74 |
| Female | 16.5-23.5<br>(18.19) | 1.98 | 1.95-2.60<br>(2.11) | 1.56 | 56.74-105.30 (72.74) | 3.65 |
| VII, Male | 24.5-30.0<br>(26.28) | 1.56 | 2.25-2.90<br>(2.44) | 1.43 | 98.35-183.74(120.52) | 2.40 |
| Female | 30.0-32.5<br>(31.15) | 1.71 | 2.65-3.40<br>(3.01) | 1.43 | 136.09-229.50 (158.29) | 2.18 |
| VIII, Male | 30.2-36.0 | 1.22 | 2.60-3.15 | 1.11 | 216.35-350.50 (261.76) | 2.17 |
|  | (32.14) |  | (2.71) |  |  |  |
| Female | 31.5-38.0<br>(34.73) | 1.11 | 3.00-3.70<br>(3.31) | 1.10 | 225.55-595.74 (382.64) | 2.42 |
| IX, Male | 42.5-51.5<br>(45.64) | 1.42 | 3.55-3.85<br>(3.68) | 1.36 | 487.35-810.30 (589.17) | 2.25 |
| Female | 45.5-52.0<br>(48.60) | 1.39 | 3.60-4.10<br>(3.81) | 1.15 | 630.35-1009.70<br>(917.46) | 2.40 |
| Mean, Male |  | 1.42 |  | 1.31 |  | 2.79 |
| Female |  | 1.44 |  | 1.32 |  | 3.00 |

### Population density and weather

Population density of *T. corallinus* varied significantly with minimum temperature and rainfall though maximum temperature showed no effect. Moreover, population density and biomass of *T. corallinus* is correlated significantly (Table 7).

**Table 7.** Result of regression analysis of different parameters.

| X-axis vs. Y-axis | R<sup>2</sup> | F | P | Equation | Significance |
| --- | --- | --- | --- | --- | --- |
| Min. temp. vs. Pop. den. | 0.3548 | 5.498 | 0.0410 | $Y = 2.018X - 17.32$ | Yes |
| Max. temp. vs. Pop. den. | 0.0277 | 0.285 | 0.6052 | $Y = 0.7175X - 2.777$ | No |
| Rainfall vs. Pop. den. | 0.3642 | 5.685 | 0.0383 | $Y = 0.0851X + 10.19$ | Yes |
| Pop. den. vs. Biomass | 0.4844 | 9.396 | 0.0119 | $Y = 89.04X + 2351$ | Yes |

## Discussion

Numbers of podous and apodous segment in each stadium overlap with the preceding or next stadium. This is a very common phenomenon in Julid millipede (Blower, 1970; Blower and Miller, 1977; David, 1982) as well as in Spirobolid (in the present study). Actually among Diplopoda, only order Polydesmida has fixed number of podous and apodous segment in each stadium.

Growth progression factor for both body length (mean 1.43) and body width (mean 1.31) is higher than a postulated value of 1.26 (Prizbram and Megusar, 1912 and Bodenheimer, 1927) or 1.28 (Horn and May, 1977 and Maiorana, 1978). Higher value for *T. corallinus* which are mainly litter feeding animal and slow mover is an affirmation of Ender’s hypothesis (1976). Mean weight progression factor of *T. corallinus* is 2.90, which is not in accordance with ‘Dyar Hutchinson Rule’ (i. e. double their weight in successive moult). Weight progression factor in female is higher than that of male and this may be due to more accumulation of egg forming reproductive tissue.

In the absence of more field data on population density of Spirobolid millipede especially *Trigoniulus* sp., some general comparison with the population of tropical species of other order will be of interest. Population density (m^-2^) of different tropical species are 4.4 (Lawrence and Samways, 2003), 15.3 – 56 (Warren and Zou, 2002), 20 – 40 (Dangerfield, 1990), 136 (Loranger, 2001), and 2300 (Bano and Krishnamoorty, 1985). In the present study it varies from 2.13 to 56.31 per m^2^. Population density of a particular species of millipedes depends on various factors like size of individual, stadia and other environmental factors i. e. availability of food, moisture and temperature. David (1987b) reported that size of millipede population is related to the quality of the humus. The better the quality, the greater will be the population size. Population density may also vary depending on elevation as observed by Alagesan and Ramanathan (2013). They reported population density of two Spirobolid (*Xenobolus carnifex* and *Aulacobolus newtoni*) in four different elevations (250m, 350m, 450m and 550m). Population density of *X. carnifex* ranges from approximately 30 /m^2^ (at 250m elevation) to 82 /m^2^ (at 450m elevation) and that of *A. newtoni* approximately 75 /m^2^ (at 250m elevation) to 162 /m^2^ (at 450m elevation).

Banerjee (1974) observed that population of *T. lumbricinus* increased from the low number in January to peak during May-July and then it declined. Here also same trend of population growth from May-August and then it declined slowly. On ecological point of view, biomass density of a species is more important than the absolute number (Critchley et al. 1979). It is expected that monthly variation of population number and biomass density should be same. But in the present study, population density is maximum in August whereas biomass density is maximum in June. Moreover, biomass density in the months of October and November is same (5605mg) though population density is more than double in October compared to that of November. This variation is due to the variation of age structure. In November, population consisted of more adult stages (which are heavier than immature) compared to October.

Millipedes have been shown to have an aggregated distribution which was often accorded with a negative binomial distribution (Banerjee, 1967a; Bhakat, 1989). Here also *T. corallinus* showed aggregated in distribution. Banerjee (1967a) calculated the aggregation of *Cylindroiulus punctatus* using k and obtained that aggregation was maximum during April to August when population density reaches maximum. He also commented that aggregation in this millipede appear to be associated with reproduction since aggregation of one sex were never found. Blower (1969) reported *Polydesmus angustus* and *P. denticulatus* are highly aggregated and the aggregations appear to be family groups. Blower (1970) measured the aggregation of *Iulus scandinavius* by using Taylor’s power law and showed that this millipede in Chesire wood are aggregated in dispersion in both soil and litter. Baker (1978c) though not measured the dispersion pattern *Ommatoiulus moreletii* but observed aggregate as beneath tussocks or underground during summer in the grassland. Mukhopadhyaya and saha (1981) showed that adults of *Orthomorpha coarctata* form aggregates in natural populations and commented that aggregate formation was due to the heterogenous distribution of resources, male attraction or egg laying habits. Bhakat (1987) also observed temporary aggregation in adults of *Streptogonopus phipsoni* between April and October. Dangerfield and Telford (1993) while studying the aggregation of a tropical millipede commented that aggregation are associated with feeding activities of immature and are not related to reproductive activity. Like other millipedes *T. corallinus* showed aggregation during the months of their peak abundance. The factor of aggregation may be many folds among them the most important factors are patchy distribution of food and soil moisture, sexual attraction in adults (Banerjee, 1967a;Dangerfield and Telford, 1993).

Banerjee (1967a) reported that in *C. punctatus* all the later stadia (from stadium VII onward) were found throughout the year but from May to September all the early stadia (stadia I to VI) were more abundant except stadium V which was found in the month of October to February. Blower and Miller (1977) also studied *C. nitidus* in a Derbyshire wood. Here all the stadia were found in the month of May and September while July and October samples mainly composed of earlier stadia. But in France, David (1982) observed that all the earlier stadia (stadia III to VIII) and adults were maximum in the month of August, September and October. Beside this, adults were also found in good number in other month of the year. In Australia, Baker (1978c) observed that stadia VIII to XII of *O. moreletii* were found throughout the year and early stadia were prevalent in the months of September to December. In *T. corallinus* early stadia (stadia II to V) were found in maximum number in the month of July, August and September but rest of the stadia was found throughout the year. So it can be concluded that *T. corallinus* follows the basic pattern of annual stadial picture as found in other millipede species.

Barlow (1958) reported that the amplitude in the seasonal population fluctuations in *Cylindroiulus friscus, Brachyiulus littoralis* and *Polydesmus denticulatus* are caused by the combined effects of rainfall and temperature, particularly of the later. Banerjee (1974) observed seasonal changes in ambient temperature and rainfall closely associated with population fluctuation of *T. lumbricinus*, though the relationship in between temperature and population density is not significant. In the present study both the abiotic factors i. e. temperature and rainfall is significantly associated with population density. Actually rainfall and ambient temperature enhance the soil and litter moisture as well as decomposition process of organic matter by microorganism and thus provide more food available within a suitable temperature range. Tracz (1984) observed that 20 species of fungi and part of various plant species decomposed by fungi are the most preferred food for *Proteroiulus fuscus*. Being ectothermic, temperature particularly minimum temperature affects the activity of millipedes. Here also millipede became surface active in the minimum temperature range 20° – 24° C.

## Acknowledgement

I am indebted to my son Dr. Soumendranath Bhakat, Lund University, Sweden for his suggestion and to all my colleagues for their inspiration.

